# Characterization of pyridomycin B reveals the formation of functional groups in antimycobacterial pyridomycin

**DOI:** 10.1101/2021.10.14.464479

**Authors:** Tingting Huang, Zihua Zhou, Maolong Wei, Lin Chen, Zhihong Xiao, Zixing Deng, Shuangjun Lin

**Affiliations:** State Key Laboratory of Microbial Metabolism, and Joint International Research Laboratory on Metabolic & Developmental Sciences, and School of Life Sciences & Biotechnology, Shanghai Jiao Tong University, Shanghai, 200240, China

**Keywords:** pyridomycin, antimycobacterial activity, biosynthesis, enoyl ester, trans ketoreductase

## Abstract

Pyridomycin, a cyclodepsipeptide with potent antimycobacterial activity, specifically inhibits the InhA enoyl reductase of *Mycobacteria tuberculosis*. Structure-activity relationship studies indicated that the enolic acid moiety in pyridomycin core system is an important pharmacophoric group and the natural configuration of the C-10 hydroxyl contributes to the bioactivity of pyridomycin. The ring structure of pyridomycin was generated by the nonribosomal peptide synthetase (NRPS) and polyketide synthase (PKS) hybrid system (PyrE-F-G). Bioinformatics analysis reveals that SDR family protein Pyr2 functions as a 3-oxoacyl ACP reductase in the pyridomycin pathway. Inactivation of *pyr2* resulted in accumulation of pyridomycin B, a new pyridomycin analogue featured with enol moiety in pyridyl alanine moiety and a saturated 3-methylvaleric acid group. The elucidated structure of pyridomycin B suggests that rather than functioning as a post-tailoring enzyme, Pyr2 catalyzes ketoreduction to form the C-10 hydroxyl group in pyridyl alanine moiety and the double bond formation of the enolic acid moiety derived from isoleucine when the intermediate assembled by PKS-NRPS machinery is still tethered to the last NRPS module, in a special energy-saving manner. Ser-His-Lys residues constitute the active site of Pyr2, which is different from the typically conserved Tyr based catalytic triad in the majority of SDRs. Site-directed mutation identified that His154 in the active site is a critical residue for pyridomycin B production. These findings will improve our understanding of the pyridomycin biosynthetic logic, identify the missing link for the double bound formation of enol ester in pyridomycin and enable creating chemical diversity of pyridomycin derivatives.

**Importance:** Tuberculosis (TB) is one of the world’s leading causes of death by infection. Recently, pyridomycin, the antituberculous natural product from Streptomyces has garnered considerable attention for being determined as a target inhibitor of InhA enoyl reductase of *Mycobacteria tuberculosis*. In this study, we report a new pyridomycin analogue from mutant HTT12, demonstrate the essential role of a previously ignored gene *pyr2* in pyridomycin biosynthetic pathway, and imply that Pyr2 functions as a *trans* ketoreductase (KR) contributing to the formation of functional groups of pyridomycin utilize a distinct catalytic mechanism. As enol moiety are important for pharmaceutical activities of pyridomycin, our work would expand the understanding the mechanism of SDR family proteins and set the stage for future bioengineering of new pyridomycin derivatives.

## Introduction

Tuberculosis (TB) remains a major global health problem, causing an estimated 10.0-million people developed TB disease in 2017 (1). With the appearance of multidrug-resistant strains of TB to all the antitubercular chemotherapies, more efficient TB control call for the development of new antimycobacterial drugs. Classic biochemical approaches, ‘-omics’ technologies including genomic- and proteomic-based approaches combined with computational chemistry strategies allow target discovery of natural products with antimycobacterial activity (2–4). Recently, reappraisal of known natural products becomes a research strategy to accelerate the development of antimycobacterial drug leads from natural products (5, 6). In *Mycobacteria tuberculosis* (*Mtb*), InhA is an NADPH-dependent enoyl-acyl carrier protein (ACP) reductase from the FASII pathway essential for synthesizing mycolic acids of cell wall (7, 8). The clinical TB drugs isoniazid and ethionamide are prodrugs that are activated by the catalase-peroxidase KatG of *Mtb* to form an adduct with NAD^+^ and then target the substrate-binding site of InhA (9). In the aim of finding new antimycobacterial compounds, pyridomycin has been identified as an InhA competitive inhibitor (10).

Pyridomycin (Figure 1A, **1**) is a cyclodepsipeptide produced by *Streptomyces pyridomyceticus* NRRL B-2517 (11). The 12-membered core ring is composed of two pyridyl groups (a N-3-hydroxypicolinyl-L-threonine and a 3-(3-pyridyl)-L-alanine), a propionic acid, and a structurally unique 2-hydroxy-3-methylpent-2-enoic acid (12). From the X-ray crystallography of pyridomycin-InhA complex, the two pyridyl groups of pyridomycin occupy the NADPH cofactor binding site of InhA, and the enolic acid (2-butan-2-ylidene) moiety protrudes into the lipid substrate binding pocket. The unique configuration that simultaneously spans both the cofactor and the substrate-binding pocket of InhA endows pyridomycin with potent activity against InhA-resistant strains of *Mtb* (10, 13).

**Figure 1.**
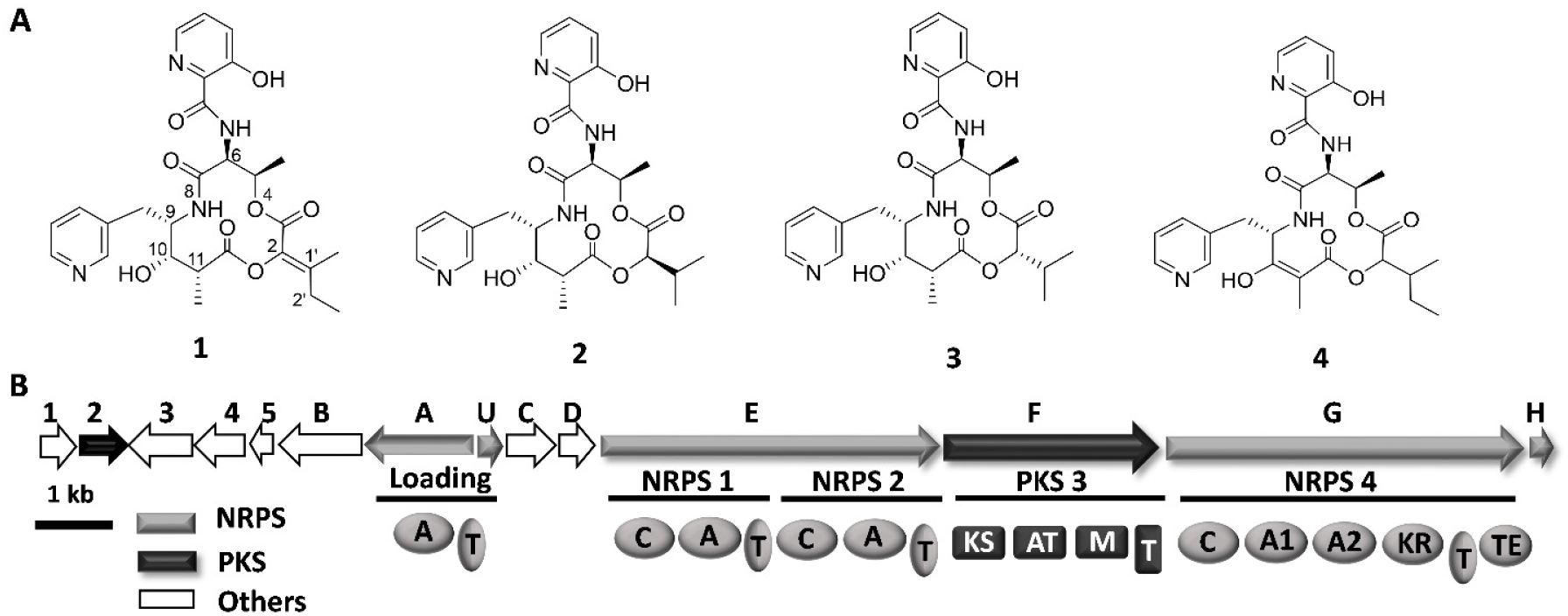
In vivo characterization of *pyr2* in pyridomycin biosynthesis. A. Structures of pyridomycin and pyridomycin B; B. Genetic organization of the pyridomycin biosynthetic gene cluster from *S. pyridomyceticus* B2517. Pyr2 is highlighted in black.

Further structure-activity relationship (SAR) investigation of chemical synthesized dihydropyridomycins (Figure 1A, **2** and **3**) revealed that substitution of the enolic acid moiety of pyridomycin by 2*R*-isopropyl moiety (**2**) made the compound less active (four-fold lower than pyridomycin) while the substitution with a 2*S*-isopropyl residue (**3**) led to diminished activity compared to pyridomycin (14). Meanwhile, the hydroxyl group on C-10 is an essential stereocenter for pyridomycin activity (15). Until now, only one total synthesis of pyridomycin has been reported (16) and the establishment of enolic acid moiety was one of the major difficulties. Motivated by the promising activity and the structural uniqueness, we set out to decipher the biosynthesis of pyridomycin both at the genetic and the biochemical level.

Nonribosomal peptide synthetases (NRPSs) and polyketide synthases (PKSs) multifunctional enzyme complexes composed of successive catalytic domains that are themselves coordinated organized into biosynthetic modules. Each module in the assembly line performs iterative chemical reactions to produce numerous natural products (17). In contrast to the co-linear manner of PKSs or NRPSs, a growing number of trans-acting units has been identified, including trans-acyltransferase (AT) for loading PKS building blocks, trans-enoylreductases (ER) for reduction of α,β-unsaturated intermediates, and trans-ketoreductase (KR) to reduce a β-ketone to a hydroxyl group (18–23). Previous biosynthetic studies showed that the pyridomycin core structure was assembled by a hybrid NRPS-PKS system consisting of PyrA, PyrU, PyrE, PyrF, and PyrG. Among them, PyrA (AMP ligase) and PyrU (peptidyl carrier protein, PCP) functioned as a free-standing adenylation (A) domain and a PCP respectively, which formed the loading module to activate and load starter unit 3-HPA (3-Hydroxy pyridine carboxylic acid) (24). PyrE, F, G encode NRPS-PKS proteins comprising four modules in total. PyrE is a typical NRPS dimodule consisting of two minimal NRPS modules (C-A-PCP) that are predicted to sequentially incorporate threonine and 3-(3-pyridyl)-L-alanine. PyrF is the sole PKS module (KS-AT-MT-ACP) and was believed to incorporate propionic acid into the pyridomycin backbone. PyrG is a special NRPS module, which contains two tandem A domains and a PKS-type ketoreductase (KR) domain (25). The second A domain in PyrG was functional for recoganization and activation of α-keto-β-methylvaleric acid (2-KVC) as the native substrate. The KR domain of PyrG further catalyzed the reduction of the 2-KVC tethered onto the PCP of PyrG to generate the PCP-tethered α-hydroxy-β-methylvaleric acid (2-HVC). A biosynthetic pathway has been proposed for the pyridomycin core structure based on these PKS/NRPS related enzymes (24). In addition, there are some other genes in the cluster with yet unassigned biosynthetic functions, which might be involved in the modification step during or after the NRPS/PKS process.

Intriguingly, according to pyridomycin structure, no KR domain was found in PyrF which was thought to be responsible for the reduction of the β-keto group to form the hydroxyl group (*). Besides, little was known about the formation of double bond of the enolic acid moiety. Close inspection of the pyridomycin gene cluster showed that *pyr2* encodes a 3-oxoacyl-acyl-carrier-protein (ACP) ketoreductase that is highly homologous to FabG from the bacterial type II fatty acid synthase. FabG is a member of the ketoacyl reductase family of proteins and is an essential enzyme for type II fatty acid biosynthesis that catalyzes an NADPH-dependent reduction of β-ketoacyl-ACP to the β-hydroxyacyl isomer, which is the first reductive step in the elongation cycle of fatty acid biosynthesis. We speculate that Pyr2, a 3-ketoacyl-ACP reductase may act as a freestanding ketoreductase in a similar manner as FabG, to reduce the β-ketoacyl intermediates tethered to PyrF-ACP (Figure 1B). By using gene inactivation and complementation, we identified *pyr2* as an essential gene for pyridomycin biosynthesis; isolation and characterization of the resulting new pyridomycin derivative from the *pyr2* mutant shed new insight into the biosynthesis of functional groups in pyridomycin. Our study not only confirmed a trans-acting NRPS/PKS specific tailoring enzyme creating chemical diversity of pyridomycin, but also provides the opportunity to explore the biosynthetic mechanism of functional moieties in pyridomycin.

## Results

### Bioinformatics analysis of Pyr2

*Pyr2*, encoding a 3-ketoacyl-acyl-carrier protein reductase, located upstream of pyridomycin NRPS-PKS genes which was originally identified as the boundary of pyridomycin gene cluster (24). Pyr2 belongs to the short-chain dehydrogenase/reductases (SDR) super family oxidoredutase with significant sequence identity (~50%) (26, 27). The amino acid sequence of Pyr2 was blast searched in microbial genome database and compared for its similarity. It reveals that Pyr2 shares sequence homology with 3-oxoacyl-ACP reductase from *Streptomyces gancidicus* (58% identity and 70% similarity) or from *Allonocardiopsis opalescens* (57% identity and 72% similarity). A database search of Pyr2 revealed homologs from various microorganisms, mostly are from high GC gram-positive strains in the phylogenetic survey. A sequence similarity network of Pyr2 with its homologs found at an e-value threshold of 10^−80^ was established (Figure 2). In the type II FAS system, 3-oxoacyl-ACP reductase catalyzes the first reduction step that leads to the conversion of 3-oxoacyl-ACP to 3-D-hydroxyacyl-ACP intermediate during the elongation cycle. Therefore, Pyr2 was considered as the most likely candidate to catalyze the C-10-keto reduction step in pyridomycin biosynthesis.

**Figure 2.**
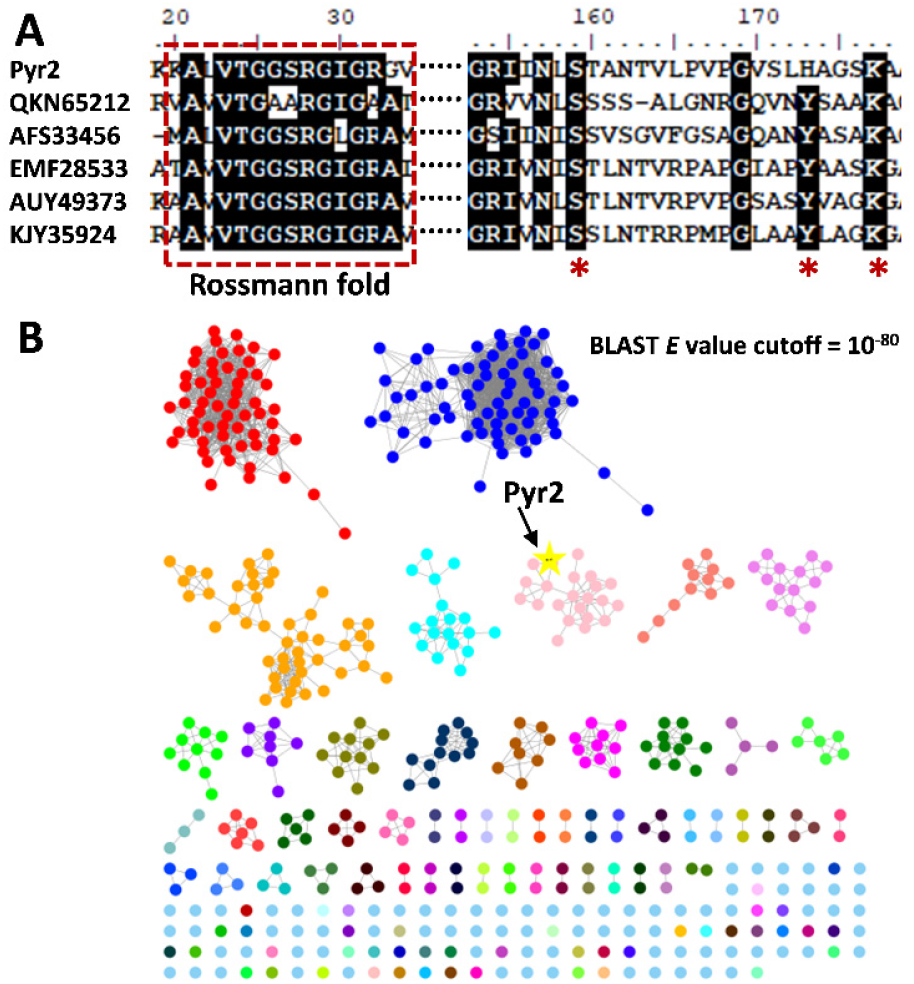
Bioinformatics analysis of Pyr2. A. Sequence alignment of selected Pyr2 homologs. The rossmann fold is highlighted in dash. The catalytic residues are shown with asterisks (*). B. A sequence similarity network (SSN) of Pyr2 homologs (sequence identity of >40%) visualized by Cytoscape.

### Inactivation and Functional Complementation of *pyr2*

To assess the necessary of *pyr2* for pyridomycin production, gene inactivation was achieved through substituting *pyr2* by the kanamycin resistance cassette using the λ_RED_-mediated PCR-targeting mutagenesis. The resultant double-crossover mutant *S. pyridomyceticus* HTT-12 was confirmed and its fermentation profile was detected by HPLC-MS (24). The mutant HTT12 retained the ability to produce a peak with the similar retention time (~16.4 min) to that of pyridomycin on HPLC profile, exhibiting a molecular ion peak at *m/z* 541 [M + H]^+^ same as that of pyridomycin (Figure 3A). The crude extracts of *S. pyridomyceticus* and HTT12 mutant were tested for antimycobacterial activity using a standard disk diffusion assay. Both of them showed similar inhibition zone on the bioassay agar plate (Figure 3B and 3C) which seems that *pyr2* is not included within the pyridomycin biosynthetic gene cluster (24).

**Figure 3.**
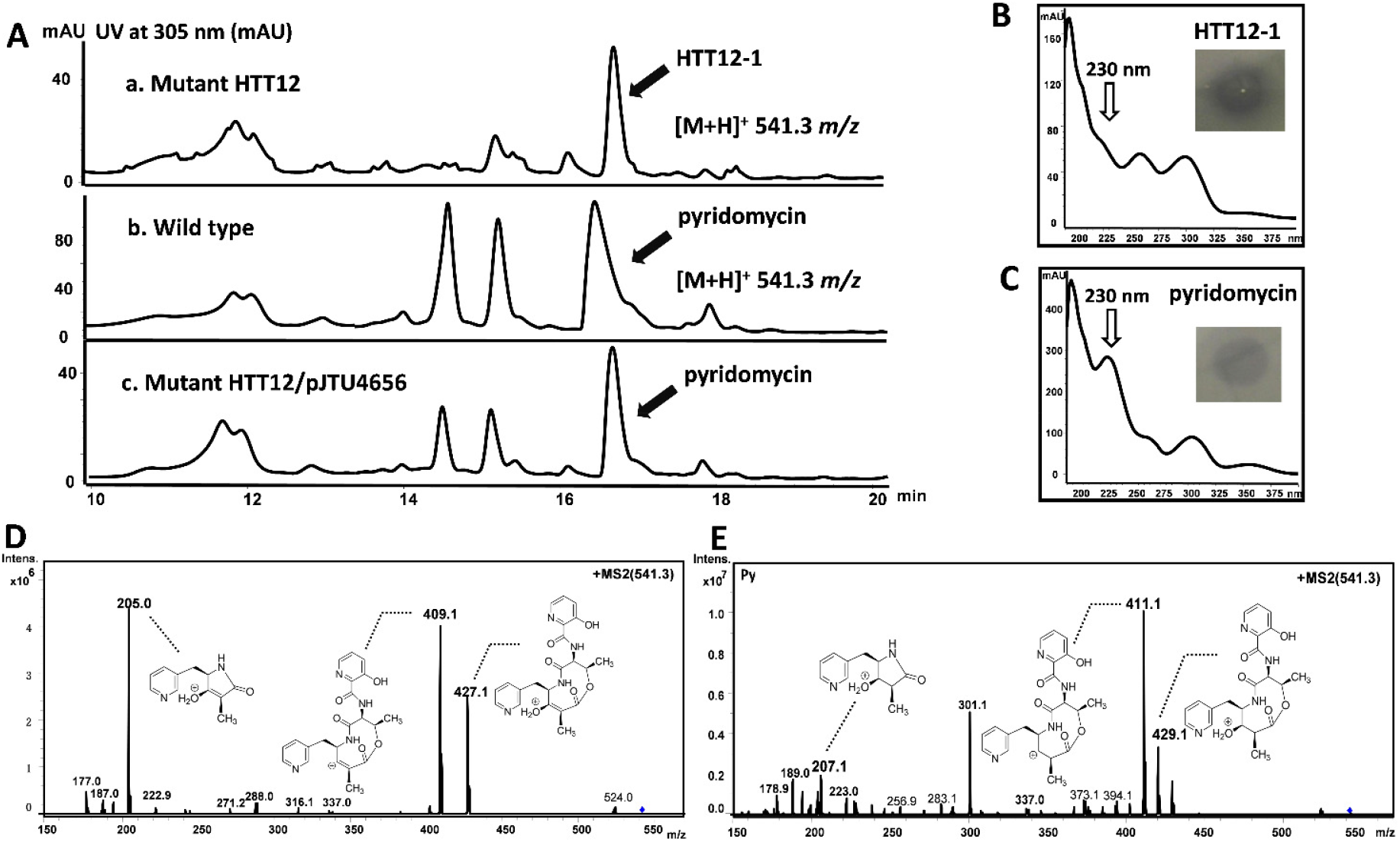
In vivo characterization of *pyr2* in pyridomycin biosynthesis. A. UV at 305 nm from HPLC analysis of metabolites from wild type *S. pyridomyceticus* B2517, HTT12 (Δ*pyr2*) and HTT12∷*pyr2*. B. Bioassay against *M. smegmatis* mc^2^155 and UV spectrum of the peak from HTT12, pyridomycin as a control (C); MS/MS spectrum of the peak from HTT12 (D) and pyridomycin (E), respectively.

Surprisingly, although the molecular weight of the product from HTT12 is as the same as that of the pyridomycin, the peak from HTT12 does not show a higher absorption shoulder at around 230 nm from the UV/Vis spectroscopy, which is slightly different from the UV/Vis spectrum feature of pyridomycin (Figure 3B and 3C). A closer inspection and comparison of the LC-MS/MS profile revealed that they had different MS/MS patterns (Figure 3E and 3F). Analysis of the peak from HTT12 in positive-ion mode showed a MS/MS fragmentation pattern (*m/z* 205.0, 409.1, and 427.1 for [M + H]^+^), presenting a −2 mass shift, compared to the fragmentation data (*m/z* 207.1, 411.1, and 429.1 for [M + H]^+^) of pyridomycin. Additionally, the two profiles also share the same ion peaks such as *m/z* at 223.0 [M + H]^+^ and 337.0 [M + H]^+^. The assignment of these fragmental ions based on the MS/MS2 data suggests that the peak in HTT12 contains a partial structure of pyridomycin and these fragments with modification on 3-(3-pyridyl)-L-alanine moiety.

To test whether the retention time shift and MS/MS pattern variation in HTT12 were actually caused by the *in vivo* deletion of *pyr2*, a complementation experiment was carried out through introduction of the integrative vector pJTU4656, a shuttle plasmid harboring a copy of *pyr2* under the control of the strong promoter *PermE*\*, into the Δ*pyr2* mutant HTT12 through conjugation. HTT12/pJTU4656 was selected in the presence of both apramycin and kanamycin. Subsequently, the culture extracts from the complementation strain were analyzed by HPLC-MS (Figure 3A, lane c). Although it was relatively less than the wild type strain, the peak in HTT12/pJTU4656 showed same MS/MS pattern as pyridomycin, indicating that pyridomycin production was restored with intact *pyr2* complementation. Taken together, the requirement of *pyr2* in the biosynthesis of pyridomycin was supported by *pyr2* inactivation and complementation. Pyr2 should play a specific role in the biosynthesis of pyridomycin.

### Isolation and structure elucidation of pyridomycin analogue from Δ*pyr2* mutant HTT12

The differences in the MS/MS patterns of the peak from Δ*pyr2* mutant HTT12 and wild type producer encouraged a large-scale fermentation to isolate the pyridomycin related compound. Following previous reported procedures (24), a 19-L culture of the mutant HTT12 was subjected for ethyl acetate extraction and subsequent column chromatography resulted in the isolation of compound **HTT12-1**.

**HTT12-1** was isolated as a white powder. The molecular formula C_27_H_32_N_4_O_8_ was established based on its HR-MS data at *m*/*z* 541.2291 (calcd. 541.2293 for [M + H]^+^), requiring 14 degrees of unsaturation (DOUs). The ^1^H NMR data of **HTT12-1** showed signals for four methyl groups [δ_H_ 0.96 (3H, t, *J* = 7.5 Hz), 1.00 (3H, d, *J* = 7.0 Hz), 1.38 (3H, d, *J* = 7.0 Hz) and 1.53 (3H, s)], four oxygenated or nitrogenated methines [δ_H_ 4.06 (1H, d, *J* = 4.5 Hz), 4.61 (1H, brs), 5.57 (1H, m) and 5.88 (1H, d, *J* = 1.5 Hz)], a 2,3-disubstituted pyridine unit [δ_H_ 8.19 (1H, d, *J* = 4.3 Hz), 7.51 (1H, dd, *J* = 8.5, 4.3 Hz) and 7.41 (1H, d, *J* = 8.5 Hz)], a *meta*-disubstituted pyridine moiety [δ_H_ 8.39 (1H, dd, *J* = 4.8, 1.3 Hz), 8.22 (1H, d, *J* = 1.3 Hz), 7.56 (1H, d, *J* = 7.7 Hz) and 7.34 (1H, d, *J* = 7.7, 4.8 Hz)] and a series of aliphatic multiplets. The ^13^C NMR spectrum, in combination with HSQC experiment resolved 27 carbon resonances consistent with four methyls, two *sp^3^* methylenes, one *sp^3^* methine, four oxygenated or nitrogenated *sp^3^* tertiary carbons, a tetrasubstituted double bond, two pyridine units, and four ester or amide carbonyl groups. As 13 of the 14 DOUs were attributed to two pyridine units, a double bond and four carbonyl groups, the remaining for DOUs required for **HTT12-1** was monocycle. The aforementioned data were similar to the pyridomycin (1), except for the absence of Δ^2 1’^ in **1** and the formation of Δ^10 11^ in **HTT12-1**(16) (Table 1, Figure S1-6).

**TABLE 1.**
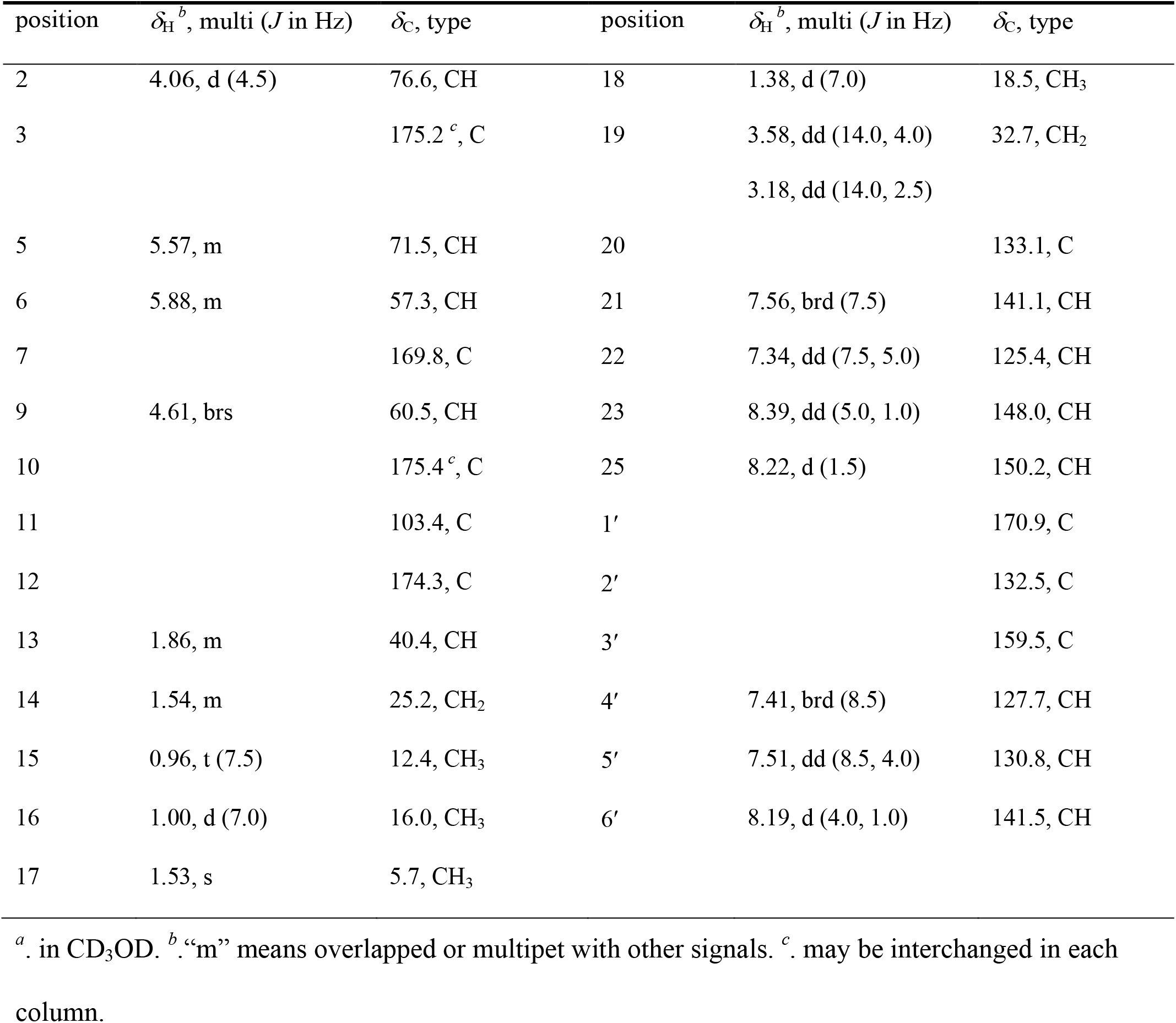
^1^H (500 MHz) and ^13^C (125 MHz) NMR data of **pyridomycin B** (*δ* in ppm).

| position | $\delta_{\text{H}}^b$, multi ($J$ in Hz) | $\delta_{\text{C}}$, type | position | $\delta_{\text{H}}^b$, multi ($J$ in Hz) | $\delta_{\text{C}}$, type |
| --- | --- | --- | --- | --- | --- |
| 2 | 4.06, d (4.5) | 76.6, CH | 18 | 1.38, d (7.0) | 18.5, CH<sub>3</sub> |
| 3 |  | 175.2 <sup>c</sup>, C | 19 | 3.58, dd (14.0, 4.0) | 32.7, CH<sub>2</sub> |
|  |  |  |  | 3.18, dd (14.0, 2.5) |  |
| 5 | 5.57, m | 71.5, CH | 20 |  | 133.1, C |
| 6 | 5.88, m | 57.3, CH | 21 | 7.56, brd (7.5) | 141.1, CH |
| 7 |  | 169.8, C | 22 | 7.34, dd (7.5, 5.0) | 125.4, CH |
| 9 | 4.61, brs | 60.5, CH | 23 | 8.39, dd (5.0, 1.0) | 148.0, CH |
| 10 |  | 175.4 <sup>c</sup>, C | 25 | 8.22, d (1.5) | 150.2, CH |
| 11 |  | 103.4, C | 1' |  | 170.9, C |
| 12 |  | 174.3, C | 2' |  | 132.5, C |
| 13 | 1.86, m | 40.4, CH | 3' |  | 159.5, C |
| 14 | 1.54, m | 25.2, CH<sub>2</sub> | 4' | 7.41, brd (8.5) | 127.7, CH |
| 15 | 0.96, t (7.5) | 12.4, CH<sub>3</sub> | 5' | 7.51, dd (8.5, 4.0) | 130.8, CH |
| 16 | 1.00, d (7.0) | 16.0, CH<sub>3</sub> | 6' | 8.19, d (4.0, 1.0) | 141.5, CH |
| 17 | 1.53, s | 5.7, CH<sub>3</sub> |  |  |  |

<sup>a</sup>. in CD<sub>3</sub>OD. <sup>b</sup>. “m” means overlapped or multipet with other signals. <sup>c</sup>. may be interchanged in each column.

The planner structure of **HTT12-1** was further established by detailed interpretation of its 2D NMR spectra. Four spin systems (C-2-C-13, C-16-C-13-C-14-C-15, C-6-C-5-C-18, and C-9-C-19) as depicted in Figure S6 were established based on the ^1^H-^1^H COSY spectrum. These fragments, ester or amide carbonyl groups, tetrasubstituted double bond, tertiary methyl, and the oxygenated or nitrogenated methines were further linked by the HMBC spectrum from H-2 to C-12, H_3_-17 to C-10/C-11/C-12, H-9 to C-10/C-20, H_2_-19 to C-10/C-21/C-25, H-5 to C-3/C-7, H-6 to C-1’/C-7, and H-4’ to C-1’, which generated a monocycle ring system. Especially, the key HMBC correlations from H-9, H_3_-17 and H_2_-19 to C-10, and from H-2 and H_3_-17 to C-12, combined with the downfield-shifted chemical shift of C-10 (δ_C_ 175.4) and upfield-shifted chemical shift of C-2 (δ_C_ 76.6), revealed the enolic hydroxy group and 1-methylpropyl group at C-10 and C-2, respectively. Thus, the gross structure of HTT12-1 was established (Figure 1). The relative configurations of HTT12-1 was assigned to be the same as pyridomycin by comparing their 1D NMR data, while the relative configurations of C-2 and C-13 were not determined. We therefore named compound **HTT12-1** as pyridomycin B (**4**) for the similar scaffold with pyridomycin.

### Determination of Pyr2 active sites

Pyridomycin B featured with enol moiety in pyridyl alanine moiety and a saturated 3-methylvaleric acid group. Inspired by the chemical structure of pyridomycin B, we reexamined the role of Pyr2. The classical SDRs family is known for sharing the canonical NAD(P)-binding motif, and this glycine-rich region plays a critical role in domain stability as well as allows access of the NAD(P) pyrophosphate. Comparison of the primary sequences of Pyr2 and its homologues revealed TGGxxGxG, the highly conserved Rossmann fold of classical SDR subfamily. In most representative 3-ketoacyl-ACP reductases, the three conserved active residues are Ser-Tyr-Lys (27). Tyr residue acts as a catalytic acid/base which forms a hydroxyl-tyrosinate ion that donates or abstracts a proton from the substrate (26). Intriguingly, alignment of Pyr2 with other canonical SDR family oxidoreductases reveals that Pyr2 contains a His instead of the usually conserved Tyr such that the active site triad is S140-H154-K158 (Figure 2).

Since the active triad S-H-K in Pyr2 is different from the conserved catalytic triad of typical SDR family proteins, we performed site-directed mutagenesis to probe the functions of the three key residues. Encouraged by the successful complementation of mutant HTT12 with Pyr2, we mutated these residues to Ala respectively using pJTU4659 as template to test *in vivo* if these Pyr2 variants retained the ability of pyridomycin B production. The resulting plasmids containing *pyr2* with point mutation were introduced into HTT12 mutant by conjugation for complementation. Pyridomycin and pyridomycin B showed some different MS/MS fragmentation patterns, we therefore chose extracted ion chromatograms (EIC) of fragment *m/z* 411 and *m/z* 409, respectively for comparison. After carefully analyzing the MS/MS profile, we found that mutating of the His154 residue to Alanine (Pyr2/H154A) cannot restore the production of pyridomycin. Both of the Pyr2/S140A and Pyr2/K158A mutant exhibit the fragment *m/z* 411, which means the complementation of pyridomycin production. Moreover, we mutated the His154 to Tyr and identified that Pyr2/H154Y mutant endow the capability of pyridomycin production (Figure 4). Such observations suggest that the catalytic triad engages in a different manner and His154 is critical for the function of Pyr2.

**Figure 4.**
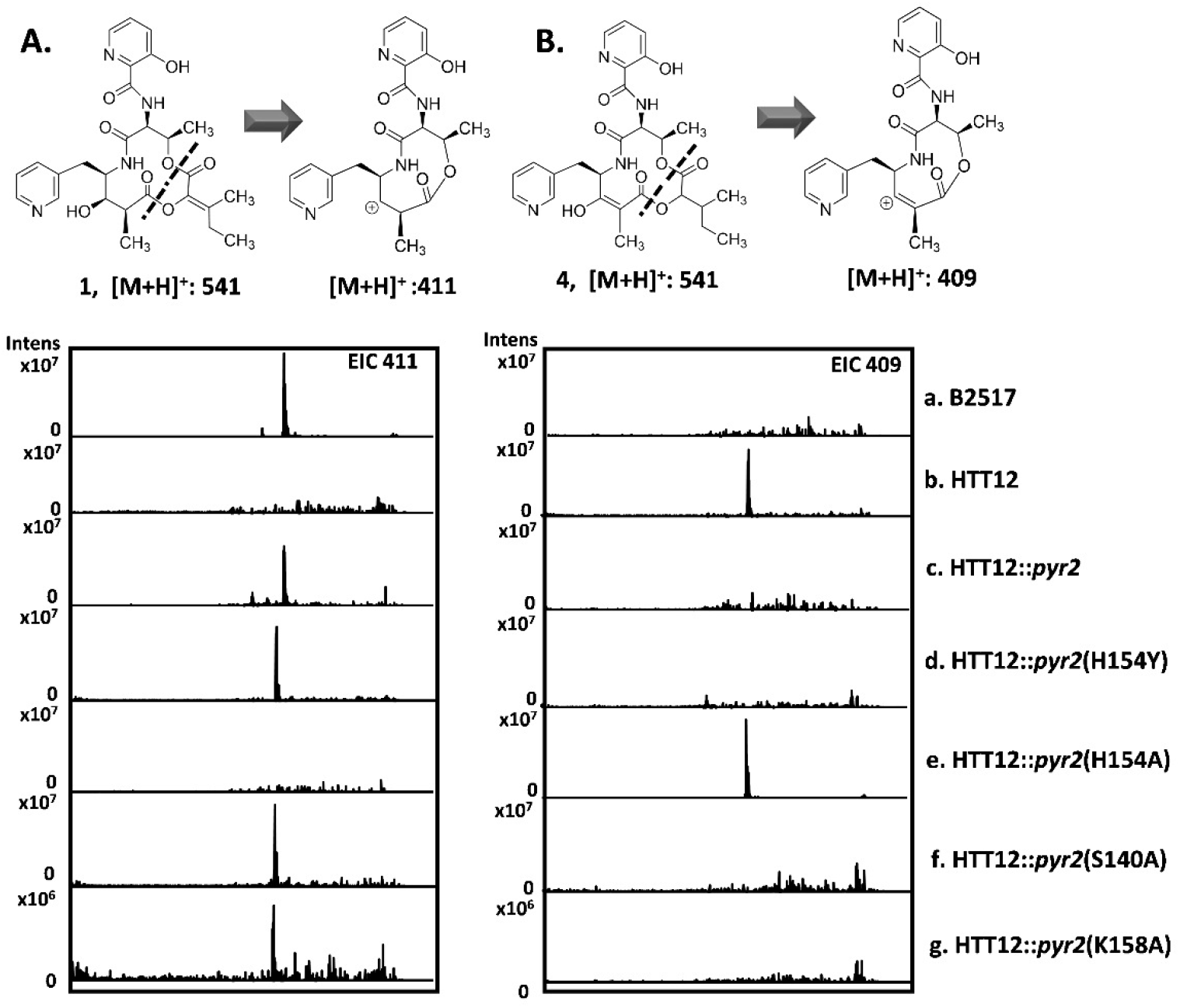
LC-MS analysis of *S. pyridomyceticus* recombinant strains. Extracted ion chromatograms (EIC, *m/z* at 409 and 411) from LC-MS analysis of metabolites from *S. pyridomyceticus* recombinant strains. The detection of each strain was individually scanned by using MS/MS fragmentation patterns of *m/z* 409 for pyridomycin and 411 for pyridomycin B.

## Discussion

Tuberculosis (TB) is one of the world’s leading causes of death. Since the discovery of pyridomycin as the antituberculous natural product over 60 years ago, it hasn’t garnered considerable attention until being demonstrated as a target inhibitor of InhA enoyl reductase of *Mtb* (28). Recently, significant progress in understanding the antituberculous mechanism inspired generation of a serious pyridomycin derivatives for SAR studies (13–15, 24). Generating pyridomycin analogues by genetic manipulation is an alternative approach based upon understanding of the biosynthetic mechanisms. In this work, we report the characterization of a new pyridomycin analogue from a *pyr2* knockout mutant HTT12, which lays a foundation for utilization of the pyridomycin pathway to develop promising antituberculous derivatives by genetic engineering.

Pyr2 is a 3-ketoacyl-ACP reductase, which belongs to the SDR superfamily. The draft genome sequence analysis of *S. pyridomyceticus* revealed the presence of additional three Pyr2 homologues (Figure S7) with >90% coverage of varying homology (>30% identities, ~52% similarities). Although the overall structure of Pyr2 and SDR proteins are highly analogous, the catalytic conformation of Pyr2 may diverge from other classical SDR family oxidoreductases as the variation of the active site triad. In addition, *Δpyr2* mutant HTT12 lost the productivity of pyridomycin, showing that these homologues could not restore the function of Pyr2. It clearly indicates that Pyr2 is pathway specific for pyridomycin biosynthesis.

Previous biosynthetic analysis revealed that among the NRPS-PKS assembly line of pyridomycin, PKS module PyrF (KS-AT-MT-ACP) incorporates the propionic acid derived unit. According to the chemical structure of final product, there is a requirement for the reduction of C-10 keto of the linear peptide-polyketide intermediate. Notably, the PKS PyrF lacks a KR domain for hydroxyl group formation, in contradiction to the presence of hydroxyl group (*) at C-10 of pyridomycin. Several examples of standalone ketoreductase from assembly-line system have been reported. For example, AntM perfroms a ketoreduction of the C-8 carbonyl of the linear ACP-bound intermediate in antimycin pathway (22). SimC7 is a ketoreductase that reduces the carbonyl to a hydroxyl group at the C-7 position in simocyclione D8 (23). It may hint that the Pyr2 contributes to the formation of the C-10 hydroxyl group. *In vivo* knockout of *pyr2* accumulated a new pyridomycin analogue, pyridomycin B. Although pyridomycin B specifically featured the enoyl group at C-10-C-11, we speculated that the enol form presented as the major species in equilibrium instead of the keto form due to the bulky aryl group of pyridomycin B at C-9 and dicarbonyl at C-10 and C-12. Isolation of pyridomycin B implies that Pyr2 functions as a *trans* ketoreductase, acts upon an acyl carrier protein-bound linear biosynthetic intermediate and catalyzes the formation of hydroxyl group on C-10 (*) prior to TE-catalyzed cyclization. We also purified Pyr2 protein and carried out *in vitro* experiment using pyridomycin B as the substrate. Unfortunately, no detectable new product in the reaction indicated that Pyr2 catalyzes a reaction on ACP or PCP-tethered substrate (data not show).

The enolic acid moiety of pyridomycin is essential for antibiotic activity, interacting with the substrate binding pocket and resulting in competitive inhibition (29). Synthesis of the enol moiety is also an obstacle in *de novo* synthesis of pyridomycin. Pyridomycin B harbors a saturated 3-methylvaleric acid group rather than an enol moiety. Based on the structure of pyridomycin B, along with the fact that there is no other dehydrogenase within this gene cluster, it seems that Pyr2 is also responsible for the double bond (C-2, **#**) formation in the isoleucine-derived moiety. We then proposed that the NAD(P) dependent enzyme Pyr2 may act as a bifunctional enzyme that abstracts the hydrogen from the α-hydroxyl-β-methylvaleric acid moiety of PyrG-tethered linear NRP/PK chain for NAD(P)H formation, and then uses the NADPH formed in situ to reduce the β-keto (*) to generate a hydroxyl group (Figure 5), suggesting an atom economy manner.

**Figure 5.**
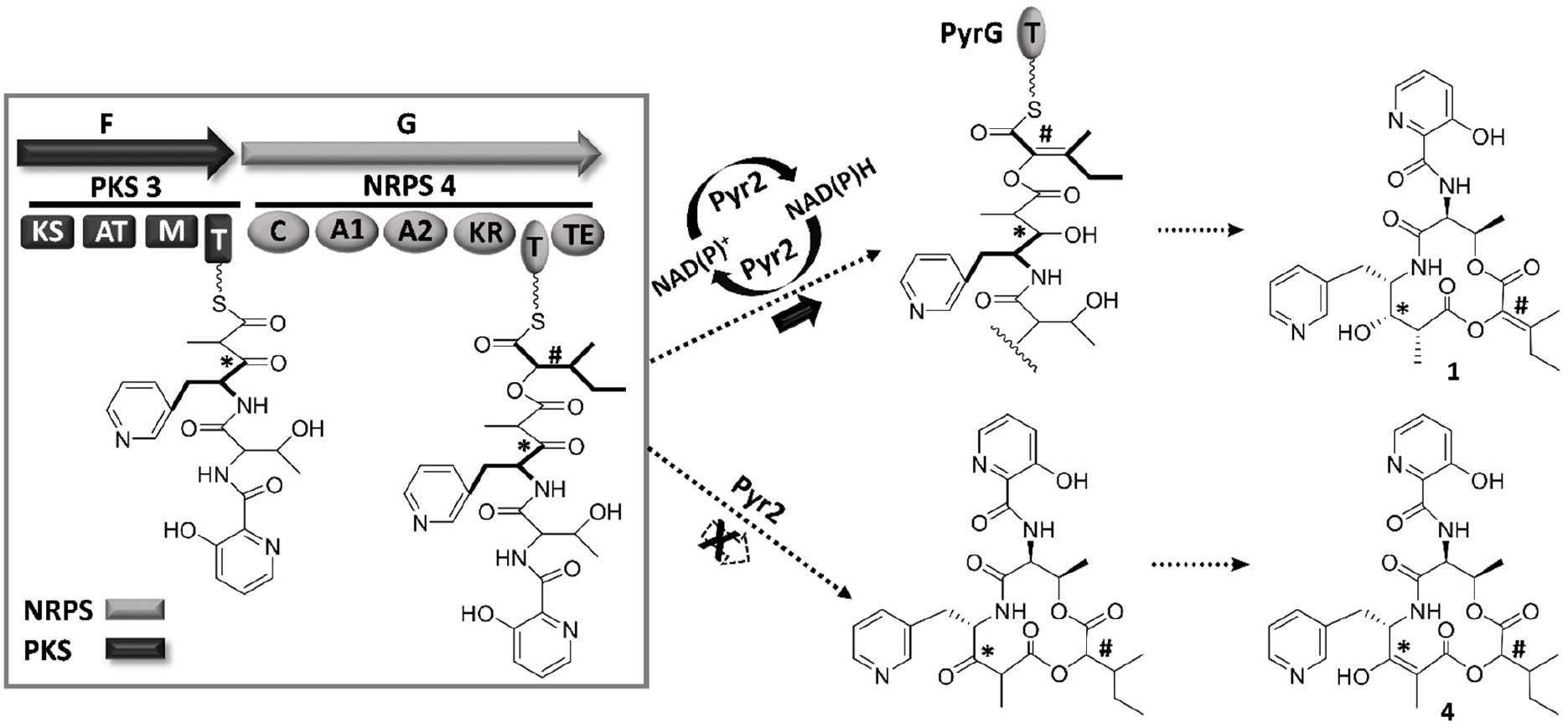
Proposed biosynthesis of pyridomycin supported by the isolation of pyridomycin B from *S. pyridomyceticus* mutant HTT12.

In summary, we confirmed that Pyr2, a 3-oxoacyl ACP reductase is essential for pyridomycin biosynthesis. The varied catalytic triad of Pyr2 suggests that the function of Pyr2 is different from the typical SDRs. The isolated pyridomycin B from the mutant Δ*pyr2* supports a biosynthetic model that Pyr2 converts the keto group to the hydroxyl group of pyridyl alanine moiety and forms the double bond of the 2-hydroxy-3-methylpent-2-enoic acid moiety using an atom economy and synergistic strategy. The enoic acid moiety is known to be an important pharmacophoric group in pyridomycin. Studies in future including functional characterization and structural elucidation of Pyr2 will lead to defining the essential motif and major active-residues of Pyr2 for interacting with the substrate, which would help to introduce other enol carboxylic acids to expand structural diversity of pyridomycin by engineering Pyr2 for antimycobacterial activity screening.

## Materials and Methods

### Bacterial strains, plasmids, and culture conditions

Bacterial strains, plasmids and oligo primers for PCR amplification used in this study are summarized in Table 2–3. PCRs were performed using rTaq (Takara biotechnology, Dalian, China) or KOD (Toyobo, Japan), and the PCR primers were synthesized by Invitrogen (Shanghai, China). Overlapping-PCR based site-directed mutagenesis was employed to introducing point mutations of *pyr2*. Restriction endonucleases were purchased from Thermo Fisher and the reactions were performed according to the manufacture’s procedures. Other common biochemical, chemical solvents and media components were purchased from standard commercial sources and used directly. *E. coli* strains were routinely grown in LB medium at 37°C and supplemented with appropriate antibiotics for plasmid maintenance. *Streptomyces pyridomyceticus* and the derived mutants were cultured in liquid tryptic soy broth (TSB) for mycelium, on solid COM medium for sporulation, on 2CM medium for conjugation and in fermentation medium (2.5% glucose, 1.5% soybean meal, 0.5% NaCl, 0.05% KCl, 0.025% MgSO_4_·7H_2_O, 0.3% K_2_HPO_4_, 0.3% Na_2_HPO_4_·12H_2_O, pH 7.2) for production of pyridomycin and pyridomycin B. *E. coli*-*Streptomyces* conjugations were performed following previously reported protocols (30).

**TABLE 2.**
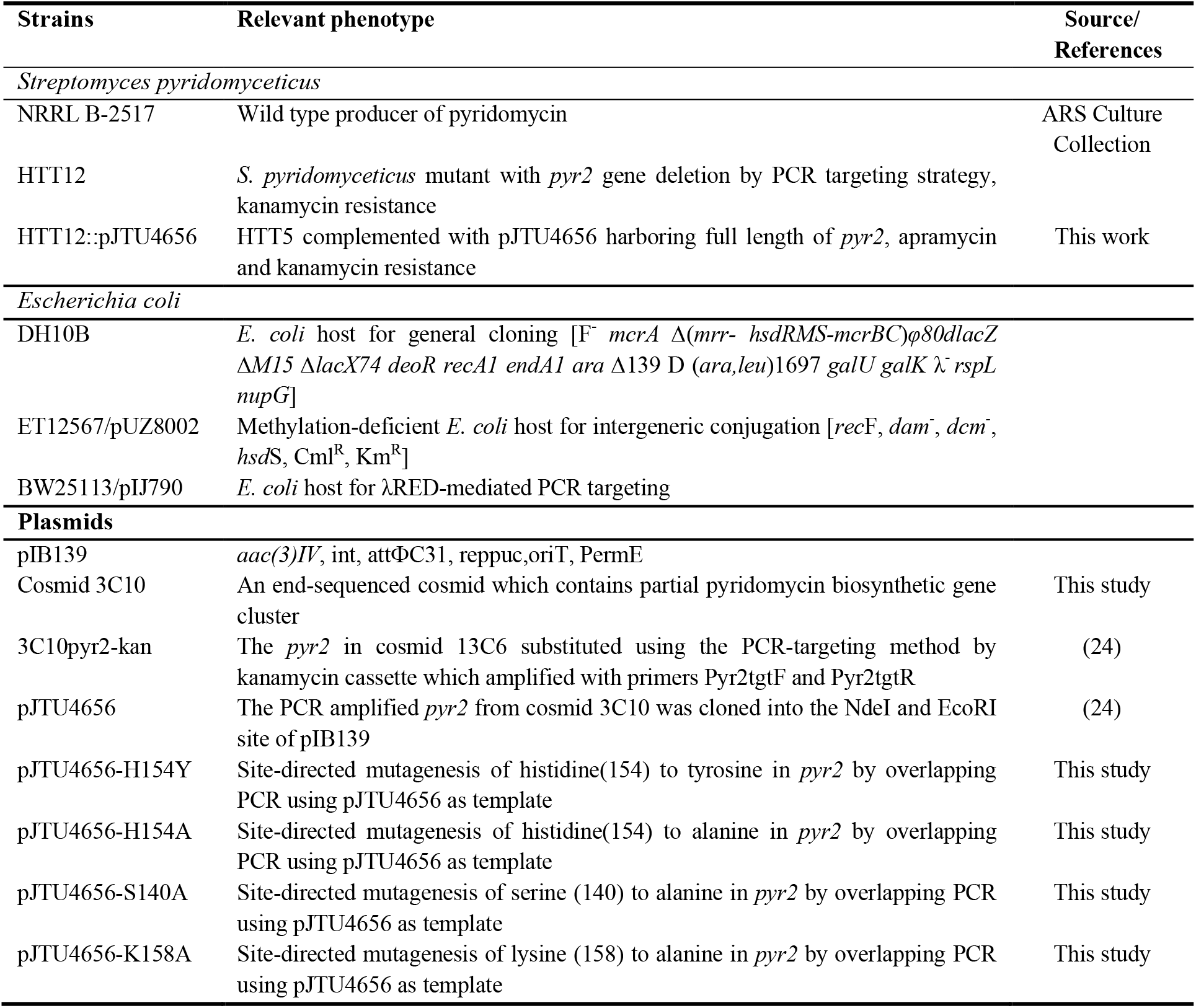
Plasmids and posmids used in this study.

**TABLE 3.**
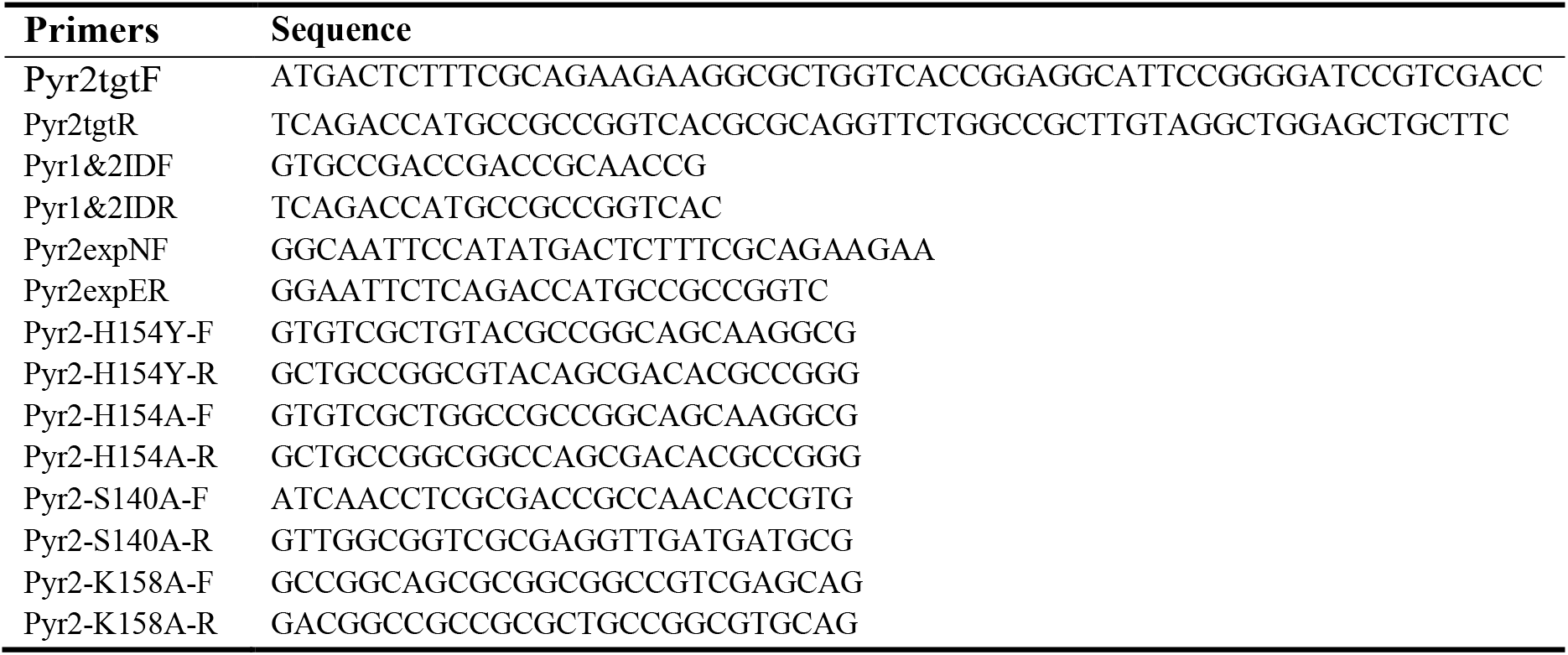
Primers used in this study.

### DNA sequencing and bioinformatics analysis

The draft genome was submitted to secondary metabolite gene cluster analysis using antiSMASH (31). A PSI-BLAST search of the NCBI nonredundant homology search was conducted on the *pyr2* and a tblastn search performed by BioEdit. The NRPS and PKS architectures were predicted by NRPS-PKS predictor (http://nrps.igs.umaryland.edu/) and confirmed against GenBank database by BLAST by manual inspection. Multiple sequence alignments and local blast analysis were performed using BioEdit Sequence Alignment Editor. The Pyr2 sequence was aligned with ClustalW using BioEdit and phylogenetic analyses were performed by MEGA-X with 1000 bootstrap replications (32). Using Pyr2 as the query, we searched for homologous proteins in the nonredundant database of Genbank and create the sequence similarity networks (SNN) using Enzyme Function Initiative (EFI) tools (33, 34). Cytoscape V6.0 was used for the SNN visualization and analysis.

### Complementation of Δ*pyr2* mutant HTT12

In *pyr2* inactivation mutant *S. pyridomyceticus* HTT12, the *pyr2* was replaced by kanamycin resistance cassette using a standard λRed-mediated PCR targeting recombination system. For the construction of the gene complementation construct, *pyr2* was amplified from *S. pyridomyceticus* genomic DNA by PCR using high-fidelity DNA polymerase and digest with *Nde*I-*Eco*RI. The resulting fragment was cloned downstream of the *PermE*\* promoter into pIB139, the *E. coli*-*Streptomyces* expression shuttle vector, digested with the corresponding restriction enzymes to yield the complementary plasmid pJTU4656 (Table S1). To achieve the point mutations in Pyr2, pJTU4656 (H154Y), pJTU4656 (H154A), pJTU4656 (S154Y), and pJTU4656 (K154Y) were constructed by site-directed mutagenesis using the primers listed in Table 2. The plasmid pJTU4656 and its derivatives carrying point mutations in *pyr2* were introduced into mutant HTT12 respectively by intergeneric conjugation as previously described. The resultant mutant strains were selected for using kanamycin resistance.

### Fermentation, production, and HPLC analysis of pyridomycin and derivatives

The Δ*pyr2* mutant HTT12 were cultured following previously reported procedures (24), with the wild type producer as a control. The seed culture was prepared by inoculating 25 mL of TSB medium in 250 mL flask with fresh spore from COM agar plate and incubating at 30°C for 24h on a rotary shaker (220 rpm). The seed culture was inoculated into a fermentation medium at 5%, and continued to incubate for 3 days. The fermentation broth was extracted three times with ethyl acetate, and the combined extract was concentrated in a vacuum evaporator to generate a solid residue. The resulting residue was dissolved in 1 mL methanol, centrifuged and filtered prior to analysis. HPLC-MS analysis was performed on a ZORBAX RX-C18 column (150×4.6 mm, 5 μm, Agilent). The column was first equilibrated with 80% solvent A (0.1% formic acid in water) and 20% solvent B (0.1% formic acid in acetonitrile). After 5 min of isocratic flow, a linear gradient was developed with a 25-min linear gradient from 20% B to 80% B at a flow rate of 0.5 mL/min and UV detection at 305 nm. The mass spectrometer was run in positive ion detection mode and set to scan between 100 and 800 *m/z*.

### Isolation and structure elucidation of pyridomycin B

Large-scale fermentation (19 L) was carried out to isolate sufficient quantities of metabolite from the Δ*pyr2* mutant HTT12 for structural elucidation. The supernatant was separated from mycelia by centrifugation (8000 rpm, 15 min), and was extracted three times with 10 L of ethyl acetate (EtOAc). The organic solvent in the combined extracts were removed under reduced pressure. The resulted crude extract (4.28 g) was adsorbed to 200 g RP-18 reverse phase resin (YMC group, Japan) and was eluted with a gradient of CH_3_OH-H_2_O (20:80, 30:70, 50:50, 70:30 and 100:0) at 3 mL/min to give 120 fractions. Each fraction was tested for alkaloid compounds by bismuth potassium iodide solution. Fractions with positive signals (fraction 69-83) were combined and concentrated to give 120 mg Fr2, further purified by a Sephadex LH-20 column using MeOH as the mobile phase at 1 mL/min to give a subfraction Fr2A (15 mg) with positive signals by TLC and UV detection. Fraction Fr2A was purified by silica gel column chromatography using gradient of H_2_O/CH_3_OH (70:30, 40:60) at 2 mL/min to yield Fr2A1 25 mg, and then subjected to semipreparative preparative HPLC (ZORBAX Eclipse XDB-C18, 9.4 × 250 mm, 5 μm, Agilent) using 0.1% formic acid in water/CH_3_OH as the mobile phase to afford pyridomycin B (25 mg).

NMR spectra were recorded on a Bruker AV-III 600 (500 MHz) spectrometer. Chemical shifts are reported in parts per million (ppm) and coupling constants *J* are given in Hertz (Hz). Structural elucidation of the isolated compounds was achieved by means of 1D (^1^H and ^13^C) and 2D (^1^H-^1^H COSY, HSQC, and HMBC) NMR spectra (Figure S1-S6) and by comparison of the spectra with published data (16). The NMR data are summarized in Table 3. Samples were dissolved in CD_3_OD. High resolution MS analysis was performed on an Agilent 1200 series LC/MSD trap system in tandem with a 6530 Accurate-Mass Q-TOF mass spectrometer with an ESI source (50-800 *m/z* mass range at positive mode).

### Antimicrobial activity assay

The antimycobacterial activities of the pyridomycin producers against indicator strain *Mycobacterium smegmatis* mc^2^155 using a standard diffusion assay. The 20 μL crude extracts of *S. pyridomyceticus* strains were dropped on paper disks and placed on the solid LB agar plates containing ~10^7^ *M. smegmatis* mc^2^155 cell. The plates were incubated overnight at 37°C and halo zone of inhibition were observed. Pyridomycin was used a positive control.

## SUPPLEMENTAL MATERIAL

Supplemental File including supplemental figure 1-7.

## ACKNOWLEDGMENTS

This work was supported by the grants from National Natural Science Foundation of China (31970053 and 32170059 to T.H., 21632007 to S.L), the National Key Research and Development Program of China (2018YFA0901900), and the Startup fund for Youngman Research at SJTU (SFYR at SJTU), respectively. We would like to thank the Instrumental Analysis Center of Shanghai Jiao Tong University and Shanghai Institute of Organic Chemistry for obtaining the NMR data.

## References

1. WHO. 2018. 2018 Global tuberculosis report. World Health Organization Press W, Geneva, Switzerland

2. Lechartier B, Rybniker J, Zumla A, Cole ST. 2014. Tuberculosis drug discovery in the post-post-genomic era. EMBO Mol Med 6:158–68.

3. Dong M, Pfeiffer B, Altmann KH. 2017. Recent developments in natural product-based drug discovery for tuberculosis. Drug Discovery Today 22:585–591.

4. Herrmann J, Rybniker J, Muller R. 2017. Novel and revisited approaches in antituberculosis drug discovery. Curr Opin Biotechnol 48:94–101.

5. Baptista R, Bhowmick S, Nash RJ, Baillie L, Mur LA. 2018. Target discovery focused approaches to overcome bottlenecks in the exploitation of antimycobacterial natural products. Future Med Chem 10:811–822.

6. Riccardi G, Pasca MR. 2014. Trends in discovery of new drugs for tuberculosis therapy. J Antibiot (Tokyo) 67:655–659.

7. Pan P, Tonge PJ. 2012. Targeting InhA, the FASII Enoyl-ACP Reductase: SAR Studies on Novel Inhibitor Scaffolds. Current Topics in Medicinal Chemistry 12:672–693.

8. Rozman K, Sosic I, Fernandez R, Young RJ, Mendoza A, Gobec S, Encinas L. 2017. A new ‘golden age’ for the antitubercular target inha. Drug Discovery Today 22:492–502.

9. Vilcheze C, Jacobs WR, Jr. 2007. The mechanism of isoniazid killing: clarity through the scope of genetics. Annu Rev Microbiol 61:35–50.

10. Hartkoorn RC, Sala C, Neres J, Pojer F, Magnet S, Mukherjee R, Uplekar S, Boy-Rottger S, Altmann KH, Cole ST. 2012. Towards a new tuberculosis drug: pyridomycin - nature’s isoniazid. Embo Molecular Medicine 4:1032–1042.

11. Maeda K, Kosaka H, Okami Y, Umezawa H. 1953. A new antibiotic, pyridomycin. J Antibiot (Tokyo) 6:140.

12. Ogawara H, Maeda K, Umezawa H. 1968. The biosynthesis of pyridomycin. I. Biochemistry 7:3296–3302.

13. Hartkoorn RC, Pojer F, Read JA, Gingell H, Neres J, Horlacher OP, Altmann KH, Cole ST. 2014. Pyridomycin bridges the NADH- and substratebinding pockets of the enoyl reductase InhA. Nature Chemical Biology 10:96–98.

14. Horlacher OP, Hartkoorn RC, Cole ST, Altmann KH. 2013. Synthesis and Antimycobacterial Activity of 2,1 ‘-Dihydropyridomycins. Acs Medicinal Chemistry Letters 4:264–268.

15. Kienle M, Eisenring P, Stoessel B, Horlacher OP, Hasler S, van Colen G, Hartkoorn RC, Vocat A, Cole ST, Altmann KH. 2020. Synthesis and Structure-Activity Relationship Studies of C2-Modified Analogs of the Antimycobacterial Natural Product Pyridomycin. J Med Chem 63:1105–1131.

16. Kinoshita M, Nakata M, Takarada K, Tatsuta K. 1989. Synthetic Studies of Pyridomycin .5. Total Synthesis of Pyridomycin. Tetrahedron Letters 30:7419–7422.

17. Wash CT, Tang Y. 2017. Natural Product Biosynthesis: Chemical Logic and Enzymatic Machinery. Royal Society of Chemistry, CPI Group (UK) Ltd, Croydon, CR04YY, UK.

18. Piel J. 2010. Biosynthesis of polyketides by trans-AT polyketide synthases. Natural Product Reports 27:996–1047.

19. Helfrich EJN, Piel J. 2016. Biosynthesis of polyketides by trans-AT polyketide synthases. Natural Product Reports 33:231–316.

20. Heneghan MN, Yakasai AA, Williams K, Kadir KA, Wasil Z, Bakeer W, Fisch KM, Bailey AM, Simpson TJ, Cox RJ, Lazarus CM. 2011. The programming role of trans-acting enoyl reductases during the biosynthesis of highly reduced fungal polyketides. Chemical Science 2:972–979.

21. Zou Y, Yin H, Kong D, Deng Z, Lin S. 2013. A trans-acting ketoreductase in biosynthesis of a symmetric polyketide dimer SIA7248. Chembiochem 14:679–683.

22. Fazal A, Hemsworth GR, Webb ME, Seipke RF. 2021. A Standalone beta-Ketoreductase Acts Concomitantly with Biosynthesis of the Antimycin Scaffold. ACS Chem Biol 16:1152–1158.

23. Schafer M, Stevenson CEM, Wilkinson B, Lawson DM, Buttner MJ. 2016. Substrate-Assisted Catalysis in Polyketide Reduction Proceeds via a Phenolate Intermediate. Cell Chem Biol 23:1091–1097.

24. Huang TT, Wang YM, Yin J, Du YH, Tao MF, Xu J, Chen WQ, Lin SJ, Deng ZX. 2011. Identification and Characterization of the Pyridomycin Biosynthetic Gene Cluster of Streptomyces pyridomyceticus NRRL B-2517. Journal of Biological Chemistry 286:20648–20657.

25. Huang TT, Li LL, Brock NL, Deng ZX, Lin SJ. 2016. Functional Characterization of PyrG, an Unusual Nonribosomal Peptide Synthetase Module from the Pyridomycin Biosynthetic Pathway. Chembiochem 17:1421–1425.

26. Kavanagh K, Jornvall H, Persson B, Oppermann U. 2008. The SDR superfamily: functional and structural diversity within a family of metabolic and regulatory enzymes. Cellular and Molecular Life Sciences 65:3895–3906.

27. Filling C, Berndt KD, Benach J, Knapp S, Prozorovski T, Nordling E, Ladenstein R, Jornvall H, Oppermann U. 2002. Critical residues for structure and catalysis in short-chain dehydrogenases/reductases. J Biol Chem 277:25677–25684.

28. Wright GD. 2012. Back to the future: a new ‘old’ lead for tuberculosis. Embo Molecular Medicine 4:1029–1031.

29. Hartkoorn RC, Pojer F, Read JA, Gingell H, Neres J, Horlacher OP, Altmann KH, Cole ST. 2014. Pyridomycin bridges the NADH- and substrate-binding pockets of the enoyl reductase InhA. Nat Chem Biol 10:96–98.

30. Kieser T, Bibb, M. J., Chater, K. F., Butter, M. J., and Hopwood, D. 2000. Practical Streptomyces Genetics: A Laboratory Manual. John Innes Foundation, Norwich, UK.

31. Blin K, Shaw S, Steinke K, Villebro R, Ziemert N, Lee SY, Medema MH, Weber T. 2019. antiSMASH 5.0: updates to the secondary metabolite genome mining pipeline. Nucleic Acids Research 47:W81–W87.

32. Kumar S, Stecher G, Li M, Knyaz C, Tamura K. 2018. MEGA X: Molecular Evolutionary Genetics Analysis across Computing Platforms. Mol Biol Evol 35:1547–1549.

33. Gerlt JA. 2017. Genomic Enzymology: Web Tools for Leveraging Protein Family Sequence-Function Space and Genome Context to Discover Novel Functions. Biochemistry 56:4293–4308.

34. Gerlt JA, Bouvier JT, Davidson DB, Imker HJ, Sadkhin B, Slater DR, Whalen KL. 2015. Enzyme Function Initiative-Enzyme Similarity Tool (EFI-EST): A web tool for generating protein sequence similarity networks. Biochim Biophys Acta 1854:1019–1037.

